# Myosin binding protein-C limits strain induced cross-bridge detachment in response to rapid stretch in cardiac and skeletal muscle

**DOI:** 10.64898/2026.03.02.709089

**Authors:** Nichlas M. Engels, Rachel L. Sadler, Michel N. Kuehn, Devin L. Nissen, Dylan Reichert, Marius Meinhold, Wolfgang A. Linke, Weikang Ma, Anthony L. Hessel, Samantha P. Harris

## Abstract

Myosin binding protein-C (MyBP-C) consists of a family of regulatory proteins expressed in sarcomeres of cardiac, fast and slow twitch skeletal muscles. The 3 MyBP-C paralogs expressed in each muscle type are encoded by separate genes but maintain a similar structure. Given the overall similarity in structure and localization of each of paralog, it is assumed that MyBP-C expressed in different muscles have similar functional effects. Here we directly tested this assumption by making use of our cut and paste approach to remove and replace N-terminal regions of MyBP-C in sarcomeres of different muscle types. We found that the different MyBP-C paralogs similarly slowed cross-bridge cycling kinetics, increased Ca^2+^ sensitivity of tension, and damped force oscillations. However, responses to a rapid stretch in actively contracting fibers, taken as indices of cross-bridge detachment and attachment kinetics, differed in each muscle type and responses depended on the presence or absence of a given paralog of MyBP-C. Altered responses to stretch were most evident for fast MyBP-C where loss of MyBP-C in psoas muscle resulted in transient responses to stretch that resembled those found in cardiomyocytes. Replacement of cardiac MyBP-C with fast MyBP-C in cardiomyocytes led to responses similar to psoas muscle. In separate X-ray diffraction experiments we also found that loss of MyBP-C in Ca^2+-^activated psoas muscle increased lattice disorder, reduced the ordering of myosin heads, and decreased thin filament length. Taken together, these results indicate that the different MyBP-C paralogs exert both common and unique effects on myosin cross-bridge kinetics.

**Significance Statement:** MyBP-C is a family of regulatory proteins found in muscle sarcomeres, where they regulate contraction and relaxation. Mutations in all MyBP-C paralogs cause disease in skeletal and cardiac muscles. We used a powerful “cut and paste” strategy to selectively remove MyBP-C from slow-twitch, fast-twitch, and cardiac muscle to show that each MyBP-C effects cross-bridge behavior similarly, though to varying degrees. Each MyBP-C had a notable effect on transient responses to rapid stretch, where MyBP-C was found to limit strain-induced cross-bridge detachment, especially in fast-twitch muscles. Strain-induced cross-bridge detachment is critical for rapid filling of the left ventricle in diastole and for sustained contraction in skeletal muscle. MyBP-C paralogs appear adapted to meet the mechanical demands of each muscle type.

## Introduction

Myosin binding protein-C (MyBP-C) consists of a family of regulatory proteins located in sarcomeres of striated muscle. Within this family, there exist three unique paralogs of MyBP-C, each encoded by a separate gene and expressed in specific fiber types. *Mybpc1* encodes the <u>s</u>low twitch skeletal paralog (sMyBP-C), *Mybpc2* encodes the fast twitch skeletal paralog (fMyBP-C), and *Mybpc3* encodes the <u>c</u>ardiac paralog (cMyBP-C)(1) The three paralogs share significant structural homology, as they all consist of a chain of 7 immunoglobulin (Ig)-like domains and 3 fibronectin type III domains termed C1-C10, a regulatory motif referred to as the M-domain, and a proline/alanine-rich domain near the N-terminus(1). Cardiac (cMyBP-C) contains an additional 8^th^ Ig-like domain at the N-terminus termed C0 (Figure 1). Each paralog is located in the C-zone of sarcomeres, where C-terminal domains C8-C10 anchor MyBP-C to the thick filament backbone and interact with myosin heads of crown 1 and 3 in relaxed muscle(2, 3). The N-terminal domains (C0-C2 for cardiac and C1-C2 for skeletal) interact with both thin and thick filaments(4–7). For example, N-terminal interactions with the thin filament shift the tropomyosin cable from the closed to the blocked state, thus promoting activation (i.e. cross-bridge formation) in the presence of Ca^2+^ or delaying relaxation as Ca^2+^ falls(8, 9). In contrast, N-terminal domain interactions with the thick filament have an inhibitory effect on myosin heads by stabilizing myosin heads in the OFF state(10, 11). While less is known regarding the middle domains (C3-C7) of MyBP-C, recent evidence suggests they play a critical auto-inhibitory role limiting the effects of N-terminal domains to activate the thin filament(12, 13). However, neither the position of the N-terminal domains nor the middle domains were resolved in recent cryo-EM reconstructions and so the structural orientation and interactions of these domains in the sarcomere are still unclear.

**Figure 1:**
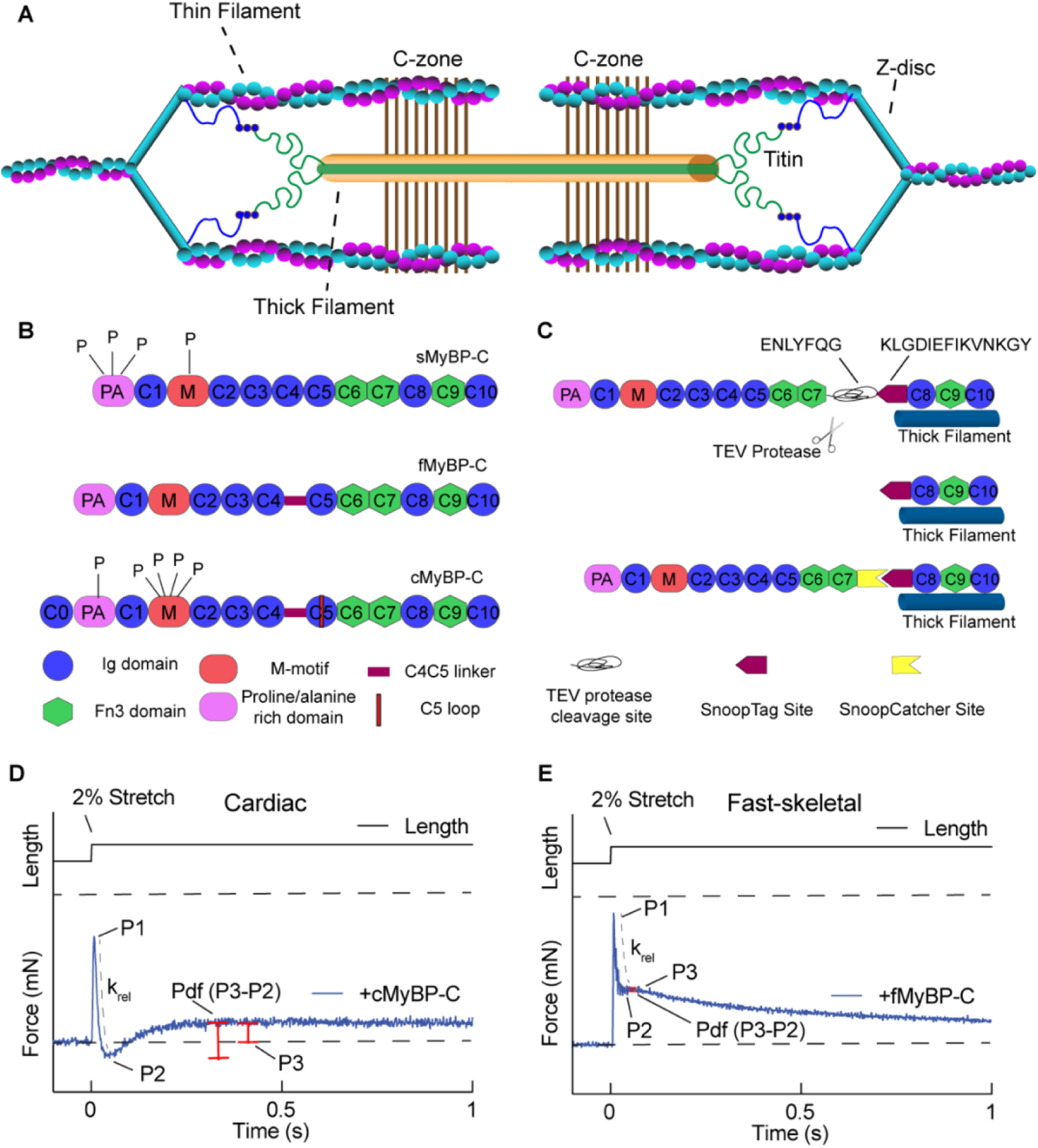
Sarcomere organization, MyBP-C structure, and utilization of the “cut-and-paste” mouse models to investigate muscle response to rapid stretch. A) Illustration of a muscle sarcomere highlighting Myosin Binding Protein-C location in the C-zone. B) Schematic of slow, fast skeletal, and cardiac Myosin Binding Protein-C paralogs. Slow and cardiac isoforms contain variable phosphorylation sites within the proline/alanine-rich as well as M-domain. Both fast and cardiac contain a linker between domains C4C5 while slow does not. Cardiac MyBP-C also contains an additional N-terminal immunoglobulin domain termed C0 as well as a 28 amino acid loop within the C5 domain. C) Schematic representation of the cut-and-paste system in fast skeletal Myosin Binding Protein-C. D) Representative force transient traces from cardiac and skeletal muscle highlighting shape differences and defining stretch activation terms for different muscle type.

As evidence for MyBP-C’s structural and regulatory roles in the sarcomere, mutations in all three MyBP-C genes lead to disease in their respective muscle types. For instance, mutations in *Mybpc3* are a leading cause of hypertrophic cardiomyopathy (HCM)(14–16), while mutations in *Mybpc1* and *Mybpc2* are linked to distal arthrogryposis (DA1) and myogenic muscle tremors(17–19). Although the three proteins share significant structural homology, sequence homology between the paralogs is low, sharing only ∼39% sequence identity between the three paralogs in human muscles(20). Additionally, sMyBP-C exists as a subfamily of multiple splice variants which likely have significant functional differences that have yet to be thoroughly characterized(21).

Here we sought to compare the relative structural and functional effects of the 3 main MyBP-C paralogs using our cut and paste approach to systematically remove and replace MyBP-C in each of the different muscle types. The approach used 3 lines of gene-edited mice, referred to as SpyC_1_, SnoopC_2_, and SpyC_3_, that allow selective removal and replacement of MyBP-C in slow twitch skeletal, fast twitch skeletal, and cardiac muscles, respectively (Figure 1)(13, 22–24). The unique strength of these mouse models is that they allow for a direct investigation of both the structural and mechanical role of MyBP-C within a single fiber type, where MyBP-C can be selectively removed and replaced in a single, healthy muscle fiber after it has been isolated from the animal. This is because all 3 mouse lines contain a tobacco etch virus (TEV) protease recognition site followed by either a short SpyTag or SnoopTag sequence between domains C7 and C8(22, 23, 25, 26) (Supplemental Figure 1). Upon treatment of permeabilized muscle fibers with TEV protease (TEVp), the endogenous domains C1-C7 in skeletal muscle and C0-C7 in cardiac muscle are selectively cleaved and eliminated from the muscle, while domains C8-C10 remain anchored to the thick filament. Following cleavage of MyBP-C, added recombinant proteins containing a cognate SpyCatcher (Sc) or SnoopCatcher(SnC) form a spontaneous covalent bond with the SpyTag or SnoopTag encoded at the N-terminus of C8, thus permitting covalent ligation (“paste”) of any desired recombinant protein *in situ* at the position of endogenous MyBP-C in the sarcomere (Figure 1).

Using these gene-edited mouse lines, we performed permeabilized muscle mechanics to measure changes in isometric force, cross-bridge cycling and stretch activation, as well as small-angle X-ray diffraction to identify structural changes to the sarcomere during isometric contraction. Here, we found that each MyBP-C paralog shares certain functional effects including sensitizing myofilaments to Ca^2+^, slowing cross-bridge cycling kinetics, and damping force oscillations at submaximal Ca^2+^ activation. However, we found that each paralog differentially regulates tension responses following a quick stretch, apparently by differentially affecting strain-induced cross-bridge detachment. In psoas muscle these changes correlated with increased thick and thin filament disorder during half-maximal isometric contraction.

## Results

### MyBP-C enhances Ca^2+^-sensitivity of tension and limits cross-bridge redevelopment rates (*k_tr_*) in cardiac muscle and fast twitch and slow twitch muscles

To determine the contribution of each MyBP-C paralog to function in the different muscle fiber types (cardiac, fast skeletal, and slow skeletal), we first investigated the effect of the different MyBP-C paralogs on steady state Ca^2+^-sensitivity of tension as well as the rate of cross-bridge cycling. Both Ca^2+-^sensitivity and crossbridge cycling, the latter of which is indicated by tension recovery from a slack-restretch maneuver, (*k_tr_)*(27) are both two parameters that are frequently altered under disease conditions. For these experiments, each MyBP-C paralog was selectively cleaved in cardiac left ventricle, fast twitch psoas, and slow twitch soleus muscles from SpyC_3_, SnoopC_2_, and SpyC_1_ mice, respectively, by incubating permeabilized muscle fibers with TEV protease (TEVp). Results shown in Figure 2 demonstrate that loss of the N’-terminal domains caused a rightward shift of the tension-pCa curve, indicating reduced Ca^2+^-sensitivity of tension such that higher [Ca^2+^] is required to achieve a given amount of force. In SpyC_3_ cardiac muscle the pCa_50_ for tension (where pCa50=-log[Ca^+^]) was shifted from 5.401 ± 0.01 prior to TEVp treatment to 5.32 ± 0.01 after treatment (n=13, *p* < 0.001) (Figure 2a). Similarly, in SnoopC_2_ fast twitch psoas muscle, the pCa_50_ was shifted from 5.63 ± 0.01 to 5.57 ± 0.01, (n=10, *p* < 0.001) (Figure 2b). In SpyC_1_ slow twitch soleus we did not observe a change in Ca^2+^-sensitivity. (5.38 ± 0.02 to 5.36 ± 0.02,n=8, n.s) (Figure 2c). However, since mouse soleus muscle expresses a near even mixture of slow and fast twitch fiber type populations with variable MyBP-C paralogs, (28) the lack of a significant difference in Ca^2+^-sensitivity of tension for soleus muscle may reflect a relatively low proportion of sMyBP-C relative to total MyBP-C. However, *k_tr_* was faster at sub-maximal levels of Ca^2+^ activation in all 3 muscle types, indicating an increase in cross-bridge cycling rates at these [Ca^2+^] (Figure 2d-f). The increased cycling suggests a change in either cross-bridge attachment or detachment rates.

**Figure 2:**
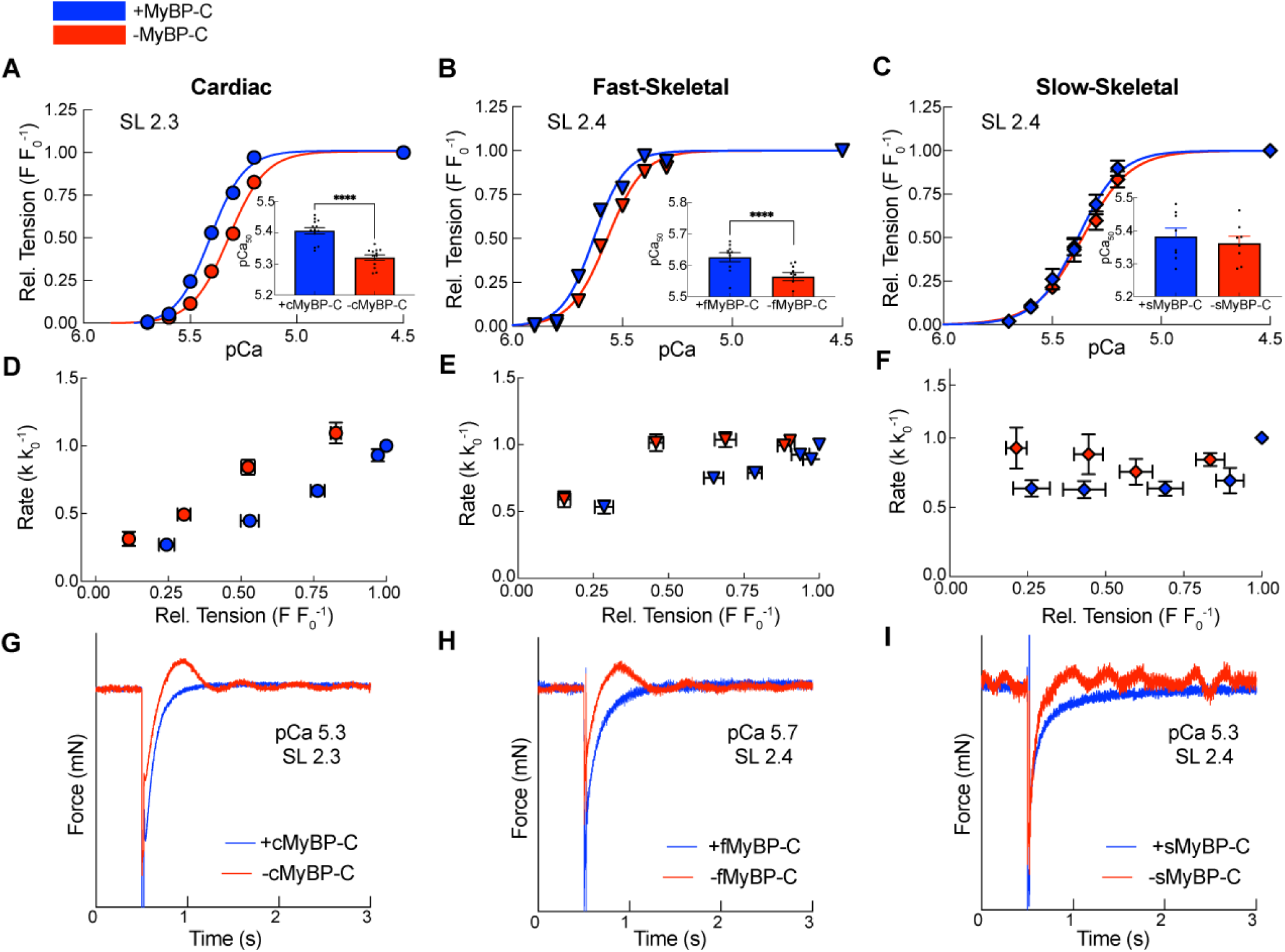
Removal of MyBP-C alters Ca^2+^-sensitivity, cross-bridge cycling rates, and produces oscillatory contractions in each muscle type. A-C) Tension-pCa curves for cardiac (A), fast skeletal (B), and slow skeletal (C) before (blue) and after (red) removal of MyBP-C. D-F) Changes in cross-bridge cycling rates in cardiac (D), fast (E), and slow skeletal (F) before and after removal of MyBP-C. G-I) Representative force traces highlighting force overshoots and tension oscillations at steady-state force in cardiac (G), fast (H), and slow skeletal (I) before and after removal of MyBP-C. Force traces only in Figure 3i (sMyBP-C) were smoothed for clearer visualization of oscillations on Graphpad Prism. Data presented as mean ± s.e.m. pCa_50_ values assessed with paired t-test. **** = p < 0.0001 in (a-c). *k_tr_* assessed with a two-way ANOVA with pCa and treatment as main effects also assessing their interaction. Individual fiber was included as a random effect. Treatment showed significance in (D-F) (D: Treatment: Prob > F = 0.0254, E: Treatment*pCa Prob > F = < 0.0033, F: Treatment Prob > F = 0.0038) A Student’s t-test or Tukey Honestly Statistically Different (HSD) post-hoc test was performed on significant main effects (p < 0.05). Dataset from: Cardiac (n=13 cardiomyocytes. N=4 mice). Psoas (n=10 fibers. N=3 mice). Soleus (n=8. N=3 +sMyBP-C. N=4 -sMyBP-C). Figure 2B, 2E, and 2H are taken from Hessel et, al. 2024 publication in Nature Communications (23).

### Loss of MyBP-C promotes the occurrence of spontaneous oscillatory contractions (SPOC) in all muscle types

Another common observation identified in each muscle type following cleavage of MyBP-C by TEVp was the appearance of spontaneous oscillatory contractions (SPOC) that became evident during Ca^2+^-activated contraction(29). The SPOC oscillations are evident as waves of contraction and relaxation that occur spontaneously, i.e., without cyclic changes in activating Ca^2+^ (see Supplemental Videos 1 and 2). Recently, the SPOC phenomenon has been proposed as an underlying mechanism of myogenic tremor in certain skeletal myopathies linked to mutations in MyBP-C(19). We have also previously reported SPOC behavior in detergent-permeabilized cardiomyocytes from SpyC_3_ left ventricles that had been treated with TEVp to remove the C0-C7 domains of cardiac MyBP-C. Importantly, the oscillations were specifically due to loss of cMyBP-C because they were eliminated by ligation of recombinant cMyBP-C(22). Here we investigated whether SPOC is a common phenomenon following removal of MyBP-C in both slow and fast twitch muscles in addition to cardiomyocytes. Figure 2g-i shows raw force traces from submaximal Ca^2+^-activated force contractions in cardiac tissue from SpyC_3_, in psoas muscles from SnoopC_2_ mice and in soleus muscle from SpyC_1_ mice before and after treatment with TEVp. Results show that removal of C1-C7 domains from both SnoopC_2_ psoas and SpyC_1_ soleus muscle consistently resulted in robust SPOC behavior which was evident as force overshoots and instabilities in the raw force traces during Ca^2+^ activation (see also Supplemental videos S1 and S2). Similar to SPOC observed in cardiomyocytes, SPOC in skeletal fibers occurred most commonly at submaximal [Ca^2+^] near the pCa_50_ for force generation. Therefore, MyBP-C paralogs appear critical in preventing SPOC from occurring in both cardiac and skeletal muscles where undamped or prolonged SPOC may contribute to muscle dysfunction in disease.

### MyBP-C paralogs differentially regulate muscle responses to rapid stretch during Ca^2+^ activation

We next investigated whether MyBP-C affects rapid tension responses following stretch in Ca^2+^ activated fibers because these responses give insights into cross-bridge detachment/attachment events as well as the delayed recovery of force referred to as SA. For these experiments, muscles were subjected to a rapid 2% stretch relative to initial muscle length followed by a 3 sec hold at the new length before returning to the initial length. The resulting transient responses to the acute stretch are typically interpreted in terms of cross-bridge behaviors(30, 31). As shown in Figure 3, the overall shape of the resulting tension transients in response to quick stretch varied significantly depending on the muscle type (i.e., responses to rapid stretch differed in cardiac, slow twitch, or fast twitch skeletal muscles).

**Figure 3:**
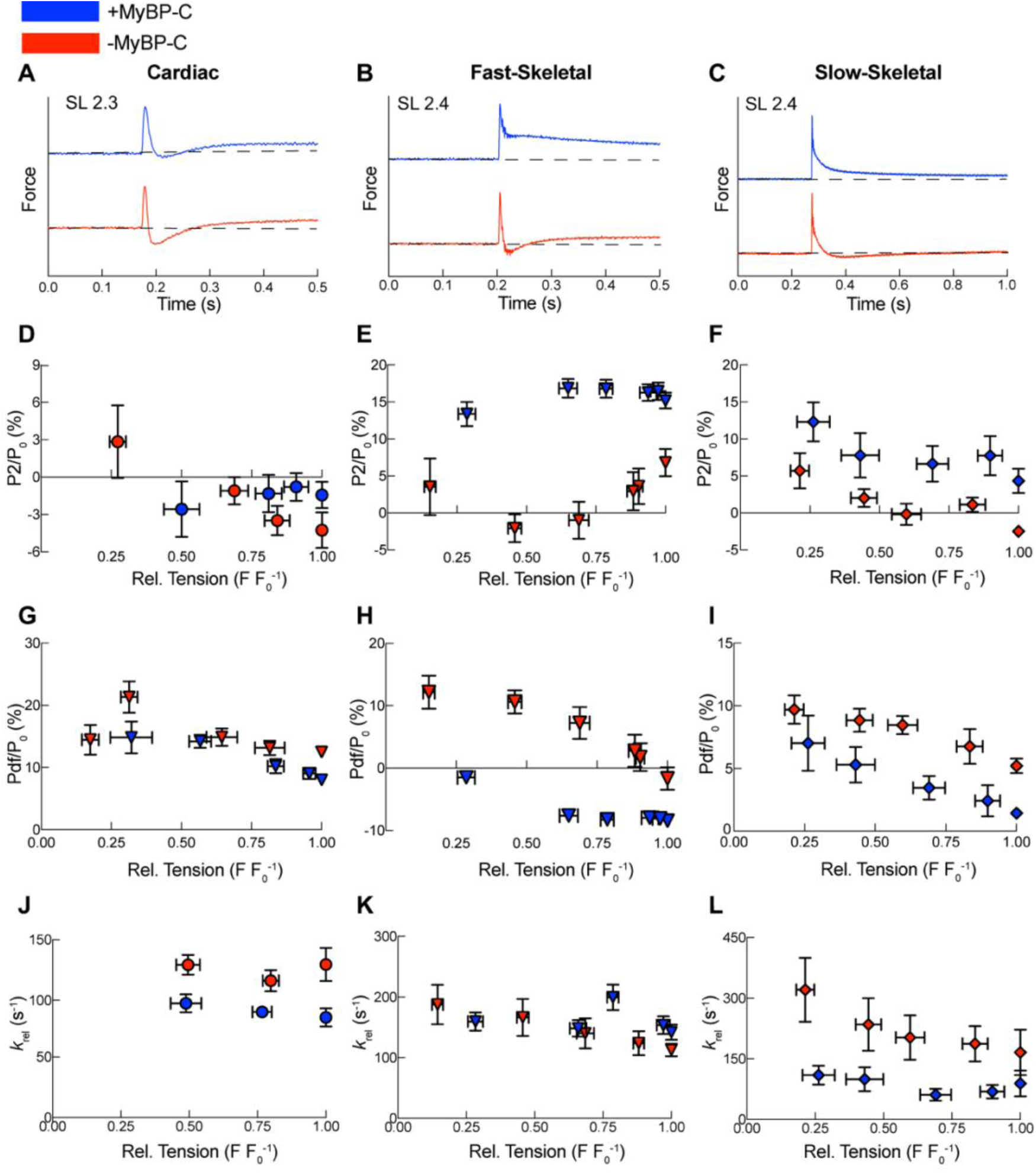
MyBP-C paralogs differentially affect tension transients following rapid stretch in each muscle type **a-c:** Representative force traces following 2% rapid stretch before (blue) and after (red)TEVp treatment of cardiac (a) fast skeletal psoas (b), and slow skeletal soleus (c). **d-f:** Mechanical measurements of the P_2_ nadir before and after TEVp treatment in cardiac (d)(pCa*Treatment Prob > F = <0.0001, fast skeletal psoas (e) (Treatment Prob > F = <0.0001, and slow skeletal soleus (f) (n.s). **g-i:** Mechanical measurements of P_df_, the amplitude of delayed force redevelopment, before and after TEVp treatment of cardiac (g)(Treatment Prob > F = 0.0001), fast skeletal psoas (h)(Treatment*pCa Prob > F = 0.0118), and slow skeletal soleus (i)(Treatment Prob > F = 0.006). **j-l:** Mechanical measurements of the rate of cross-bridge relaxation (*k_rel_*) following rapid 2% stretch before and after TEVp treatment of cardiac (j)(Prob > F = <0.0001), fast skeletal psoas (k)(Treatment: Prob > F = 0.0272), and slow skeletal soleus (l)(n.s). Data presented as mean ± s.e.m. A repeated measures ANOVA design was performed throughout with pCa, treatment, and their interaction as main effects. A random effect was included for each individual fiber. A Student’s t-test or Tukey’s HSD post-hoc test was performed on significant main effects (p < 0.05). Dataset: Cardiac from (n=13 cardiomyocytes. N=4 mice). Psoas (n=10 fibers. N=3 mice). Soleus (n=8. N=3 +sMyBP-C. N=4 -sMyBP-C). Further detail in supplement.

Figure 1 shows representative traces from cardiac muscle and psoas muscle that illustrate the 3 typical phases of the transient force responses to stretch. In cardiac muscle following rapid stretch there is a rapid increase in force (P_1_) that coincides with and is proportional to the initial stretch/increase in length. P_1_ is related to both strain on bound cross-bridges as well as strain on elastic elements within the sarcomere, such as titin(32). Following the P_1_ peak, the force drops rapidly in such a way that the rate of the decrease in tension (K*_rel_*) is related to the kinetics of cross-bridge detachment. Next, the nadir of the tension decrease is termed P_2_ and typically approaches or drops below the pre-stretch steady state tension (33). Following P_2_, there is a delayed rise in tension where tension increases to a new steady-state value that is greater than the pre-stretch steady state force (P_3_). This rise to a new steady state force (P_3_) termed SA (33–35), is most prominent in muscles that do oscillatory work such as some insect flight muscles and vertebrate cardiac muscle, whereas it is typically less prominent and shorter lived in skeletal muscle(34, 36). In fast twitch skeletal muscle, the rapid increase to P1 is similar to cardiac muscle. However, there is a notable difference following the initial relaxation between cardiac and fast twitch skeletal muscle, where P_2_ typically remains greater than the steady state force prior to stretch(34). Next, instead of a delayed rise in tension to a new steady state (P_3_), the force briefly plateaus before slowly decreasing to a steady-state. Since the fast twitch muscle tension transient typically lacks a delayed regeneration of force, the SA response is considered minimal in fast twitch skeletal muscle.

Here, we found that the tension responses to rapid stretch were affected following removal of MyBP-C in each muscle, albeit, to varying degrees. A representative trace for each muscle type before and after removal of MyBP-C is shown in Figure 3. Results show that removal of cMyBP-C in cardiomyocytes had overall modest effects on tension transients, i.e. loss of cMyBP-C slightly reduced the P_2_ nadir (by ∼4% at pCa 4.5 and 5.2), increased the P_df_ amplitude (P_3_-P_2_) by ∼4% at pCa_50_, and increased *k_rel_*, the rate of relaxation following the P_1_ peak (97.56 ± 7.37 to 129.37 ± 8.20 at pCa_50_) (Fig. 3d,g,j). However, effects of removing fMyBP-C in psoas muscle were much more profound where tension transients following loss of fMyBP-C (Figure 3b) now closely resembled a typical transient response seen in cardiac muscle. That is, removal of fMyBP-C significantly decreased the P_2_ nadir below the pre-stretch steady state and increased the P_df_ amplitude. K_rel_ was also significantly increased at sarcomere length (SL) 3.0 (Supplementary Figure 2), although K*_rel_* remained unchanged at SL 2.4 possibly due to a lower rate of data acquisition for this data set (Figure 3e,h,k). Most strikingly, the acute removal of fMyBP-C resulted in the appearance of a new delayed phase of force regeneration (phase 3) that is usually minimal or absent in fast twitch fibers. This was primarily due to the decrease in P_2_ because the P_3_ amplitudes remained unchanged in cardiac and skeletal muscle. However, the P3 does appear slightly increased in the presence of fMyBP-C (Supplemental Figure 3). In slow twitch soleus muscle, similar to cardiac muscle, the overall shape of the transient was not dramatically changed following loss of sMyBP-C, but both the P_2_ nadir decreased further beyond the pre-stretch steady state, P_df_ amplitude increased, and K*_rel_* rates were also increased (Figure 3f,i,l). Taken together, loss of fMyBP-C conferred SA-like properties to fast skeletal fibers, suggesting that canonical cross-bridge tension transient responses to stretch are modulated by MyBP-C paralogs in the different fiber types.

### Skeletal muscle MyBP-C paralogs alter tension transient properties in cardiac muscle

To directly test the extent to which differences in transient tension responses to stretch are due to MyBP-C paralog expression in different muscle types, we next assessed whether fMyBP-C and sMyBP-C could confer skeletal-like responses to stretch in cardiomyocytes. We chose cardiac muscle as the background for paralog exchanges in this set of experiments because we were able to remove nearly all endogenous cMyBP-C following TEV protease treatment in cardiac myocytes compared with the more limited removal of skeletal muscle paralogs from either psoas or soleus muscles that express a mixture of skeletal MyBP-C paralogs (22). Additionally, because mouse hearts contain primarily a single myosin isoform (α-myosin)(37), other potentially confounding effects due to variable myosin isoform expression in different skeletal muscle fiber types are avoided(38). Therefore, to directly assess effects of the different MyBP-C paralogs, cardiomyocytes from SpyC_3_ mice were first treated with TEVp to eliminate endogenous cMyBP-C and then cMyBP-C was replaced *in situ* with either recombinant cMyBP-C, fMyBP-C or sMyBP-C via SpyCatcher encoded at their C’-terminal ends. As shown in Figure 5a, recombinant cMyBP-C was first added as control. Ligation of recombinant cardiac C0C7Sc did not alter the overall shape of the tension transient compared to before TEVp treatment, although the P_2_ value tended to be higher. This could be due to the lack of post-translational modifications (e.g., phosphorylation) of the recombinant C0C7Sc compared to the endogenous cMyBP-C prior to TEVp treatment since phosphorylation of cMyBP-C reduces P_2_(39). Ligation of cardiac C0C7Sc also restored Ca^2+^-sensitivity of tension (pCa_50_: Pre: 5.41 ± 0.01, Post: 5.32 ± 0.01, cC0C7Sc: 5.39 ± 0.02) and reduced *k_tr_* rates to values before treatment with TEVp (Figure 4a,d). Similar effects were observed for ligation of fast C1C7Sc, where Ca^2+^-sensitivity was restored (pCa_50_: Pre:5.54 ± 0.02, Post:5.44 ± 0.01, fC1C7Sc: 5.54 ± 0.01) (Fig. 4.b,e). However, after ligation of fC1C7Sc transient responses to stretch in cardiac muscle were altered such that they more closely resembled those typically seen in fast twitch skeletal fibers instead of those seen in cardiomyocytes. That is, fC1C7Sc significantly elevated the P_2_ amplitude above the pre-stretch steady state force and nearly eliminated the rise in P_3_ force (SA), leading to negative P_df_ values (Figure 5e,h). Additionally, fC1C7Sc decreased the phase 2 relaxation rate K_rel_, trending to be slightly slower than endogenous cMyBP-C. (Fig. 5k). Taken together, these experiments demonstrate that removal of fMyBP-C from psoas muscle altered the tension transient to resemble cardiac muscle, while addition of recombinant fC1C7Sc to cardiac muscle altered the tension transient to resemble fast twitch skeletal muscle.

**Figure 4:**
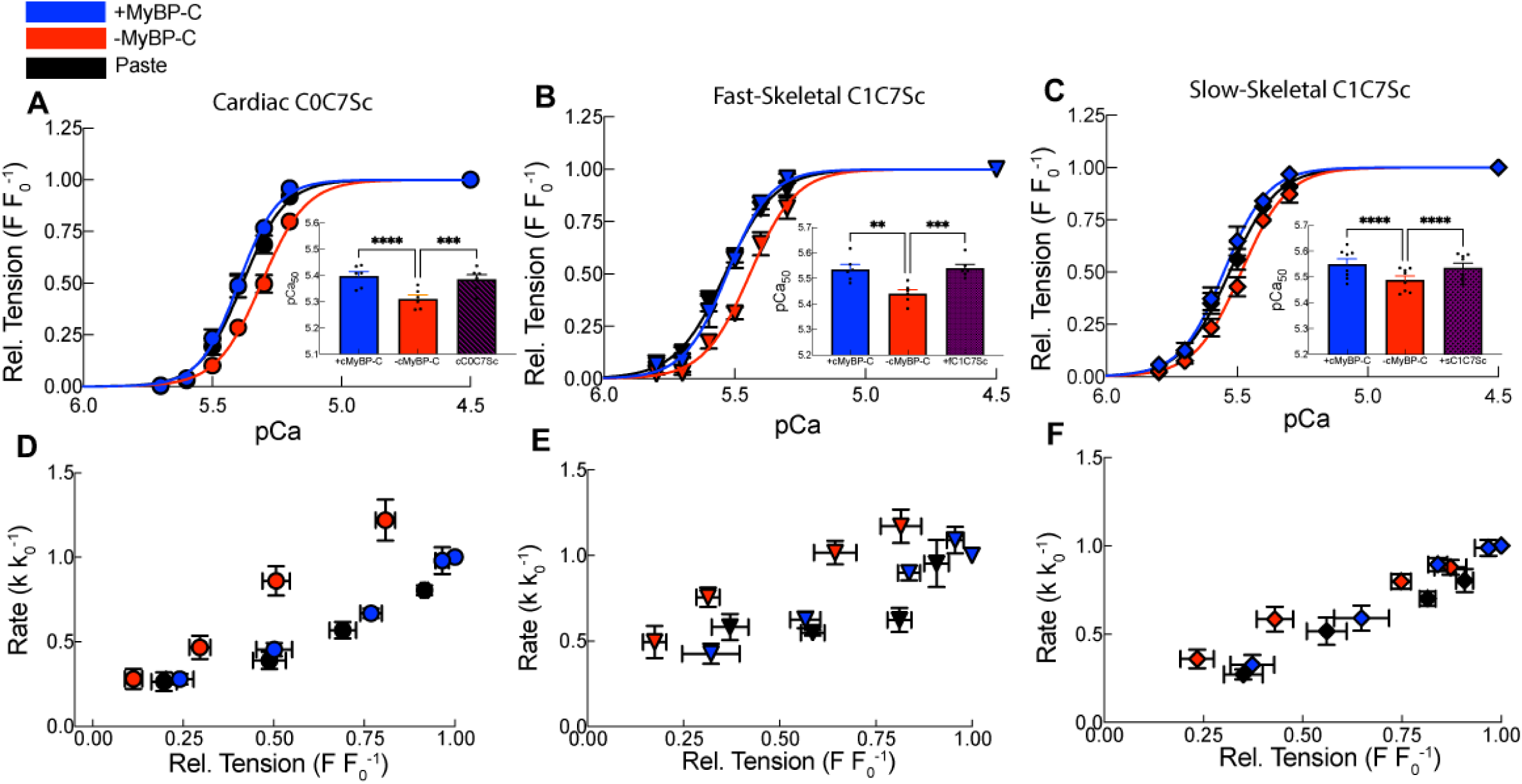
Cut-and-paste experiments performed in detergent permeabilized cardiac muscle to assess Ca^2+^-sensitivity and *k_tr_* after *in situ* replacement of recombinant cardiac, fast skeletal, and slow skeletal MyBP-C. **a-c:** Relative tension-pCa with inset containing average pCa_50_ following each curve for cC0C7Sc(a), fC1C7Sc (b), and sC1C7Sc (c). ** = p = <0.0001, *** p < 0.0005, **** p = <0.005 **d-f:** Changes in *k_tr_* before and after removal of endogenous cMyBP-C and following paste of cC0C7Sc (d) (pCa*Treatment Prob > F = 0.0046), fC1C7Sc (e) (Treatment Prob > F = 0.0061) and sC1C7Sc (f) (Treatment Prob > F = 0.0046). Statistics throughout are an ANOVA design with pCa, treatment, and their interaction as main effects, as well as an individual fiber random effect. Statistically significant differences were assessed using a Tukey’s HSD post-hoc test on statistically significant differences (p < 0.05). Dataset: cC0C7Sc (n=7, N=3), fC1C7Sc (n=6, N=5), sC1C7Sc (n=8, N=6).

**Figure 5:**
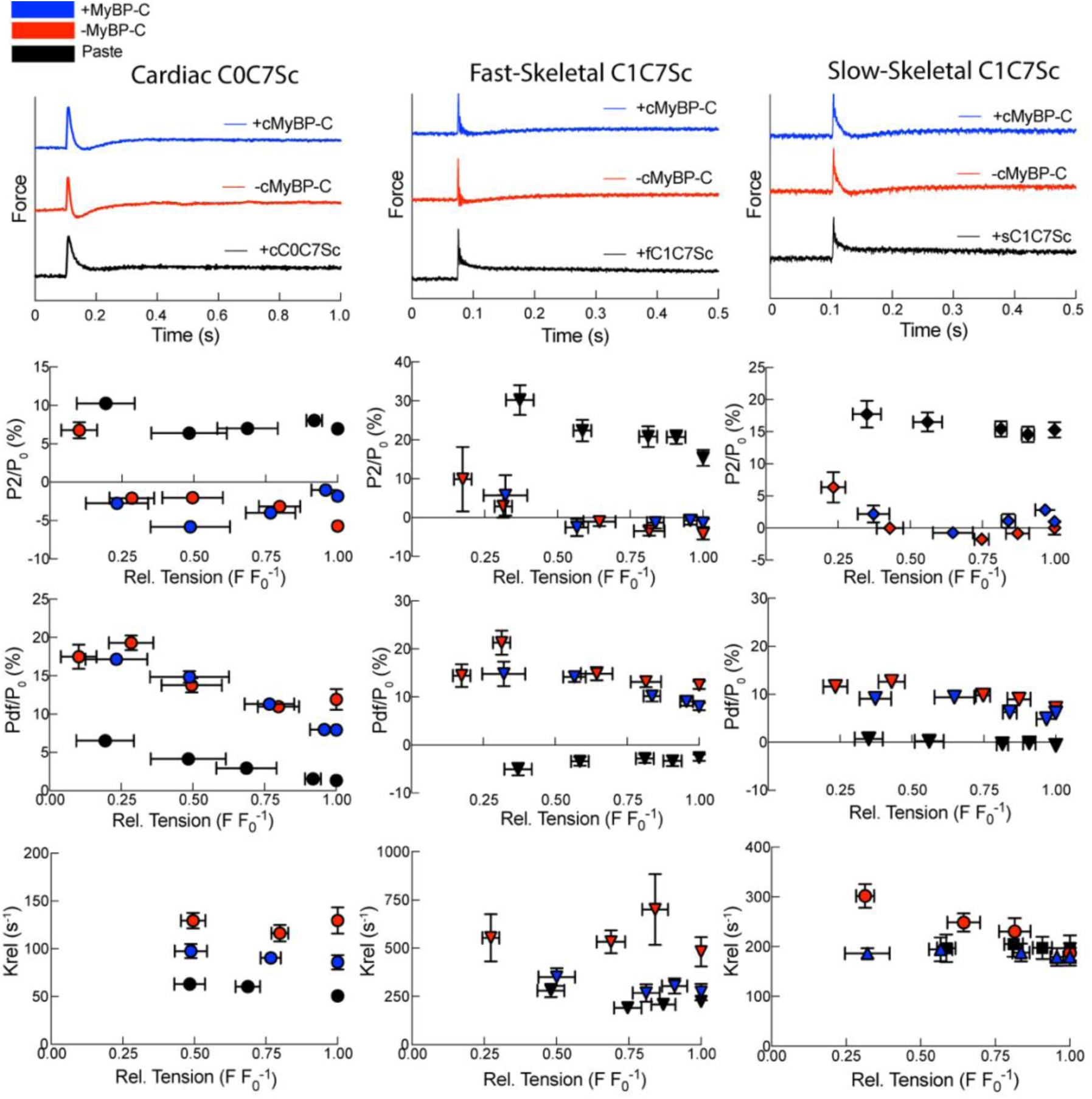
Amplitudes of strain-induced cross-bridge detachment (P_2_), stretch activation (P_df_), and cross-bridge relaxation kinetics (*k_rel_*) in cardiac muscle following *in situ* replacement with cC0C7Sc, fC1C7Sc, and sC1C7Sc. **a-c:** Representative force traces of cardiac tension transients before (blue) and after (red) removal of endogenous cMyBP-C, and following replacement (black) with recombinant cC0C7Sc (a) fC1C7Sc (b), and sC1C7Sc (c). **d-f:** Amplitude of strain induced cross-bridge detachment (P_2_) before and after removal of endogenous cMyBP-C and after replacement with cC0C7Sc (d) (pCa*Treatment*pCa Prob > F = <0.0001), fC1C7Sc (e) (Treatment Prob > F = <0.0001), and sC1C7Sc (f))(Treatment*pCa Prob > F = 0.0234) **g-i:** Amplitude of stretch activation (P_df_) before and after removal of endogenous cMyBP-C and after *in situ* replacement with cC0C7Sc (g)(Treatment*pCa Prob > F = 0.0029), fC1C7Sc (h)(Treatment*pCa Prob > F = 0.0076), and sC1C7Sc (i)(Treatment*pCa Prob>F = 0.0181. **j-l:** Rates of stretch induced cross-bridge relaxation (*k_rel_*) before and after removal of endogenous cMyBP-C and after *in situ* replacement with cC0C7Sc (j)(Treatment Prob > F = <0.0001), fC1C7Sc (k)(Treatment Prob > F = <0.0001), and sC1C7Sc (l)(Treatment*pCa Prob > F = 0.0076). Statistics throughout are an ANOVA design with pCa, treatment, and their interaction as main effects, including an individual fiber random effect. A post-hoc Tukey’s HSD test was performed on statistically different main effects (p < 0.05). Dataset: cC0C7Sc (n=7, N=3), fC1C7Sc (n=6, N=5), sC1C7Sc (n=8, N=6).

Similar results were seen following replacement with sC1C7Sc, which was able to restore Ca^2+^-sensitivity of tension (pCa_50_: Pre 5.55 ± 0.02. Post: 5.49 ± 0.01, sC1C7Sc: 5.54 ± 0.02) and reduce k*_tr_* rates to a similar extent as addition of cMyBP-C (Figure 4c,f). Therefore, both recombinant fMyBP-C and sMyBP-C are sufficient to restore typical interactions of endogenous cMyBP-C in cardiac muscle. sC1C7Sc also raised the P_2_ nadir to a level above pre-stretch steady state and nearly eliminated P_df_ amplitude of the tension transient, thus significantly blunting the SA response typically seen in cardiac muscle (Fig. 5f,i). Taken together, these results demonstrate that skeletal muscle paralogs of MyBP-C reduce canonical SA responses in cardiac muscle and instead confer skeletal-like responses to rapid stretch.

### Structural changes following loss of fMyBP-C determined by X-ray diffraction

The startling result that fMyBP-C cleavage in psoas leads to cardiac-like SA properties encouraged us to identify underlying sarcomere-level structural changes. We therefore used the small-angle X-ray diffraction technique on permeabilized psoas fibers from SnoopC_2_ mice under Ca^2+^-activated conditions to determine structural changes in the sarcomere before and after cleavage of fMyBP-C. In particular, we sought to determine whether fMyBP-C contributed to changes in lattice spacing, lattice order, and mass distributions on the thick or thin filaments that could potentially account for the change in transient force behaviors described above. X-ray reflections and their relationship to sarcomere structures can be visualized in the cross-sectional diagram of the sarcomere (Fig. 6a,b,c).

**Figure 6:**
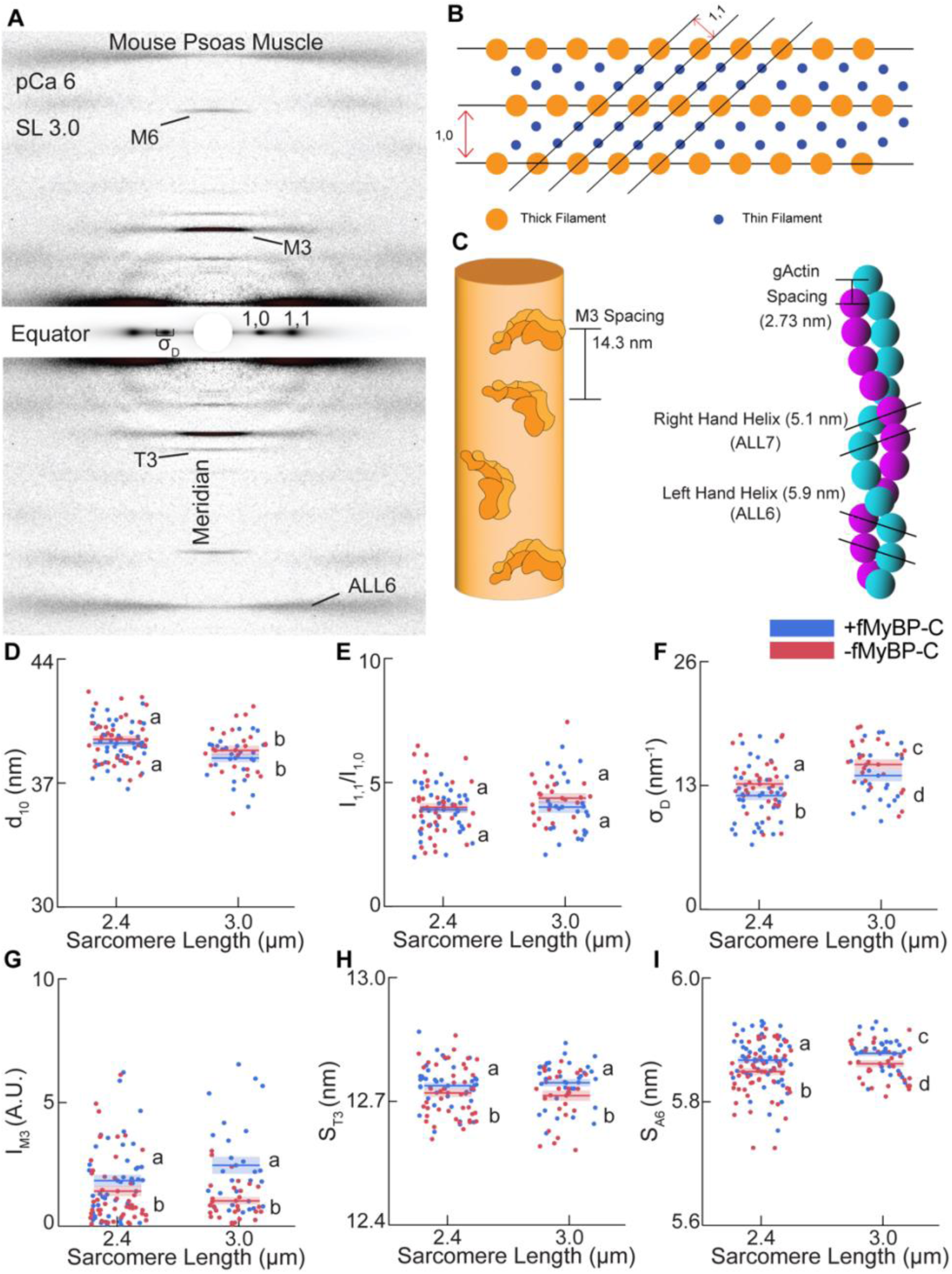
Equatorial and meridional reflections of isometrically contracting psoas Muscle in presence and absence of fMyBP-C. **a)** Representative X-ray diffraction image at SL 3.0 and pCa 6.0 highlighting meridional reflections (M3, T3, M6, A6) assessed in this study. **b)** Graphic showing a cross-sectional view of the sarcomere lattice as well as important equatorial reflections. **c) Left:** Schematic of the quasi-helical arrangement of myosin heads on the thick filament backbone, where the M3 reflection is highlighted as difference in space between individual myosin crowns. **Right:** Schematic of the thin filament, highlighting both the left-handed and right-handed helix of the actin monomers and their spacings. **d)** Lattice spacing of permeabilized psoas muscle at SL 2.4 and SL 3.0 before and after removal of fMyBP-C (Treatment: Prob > F 0.1494) **e)** I_11_/I_10_ intensity ratio of permeabilized psoas muscle at SL 2.4 and 30 before and after removal of fMyBP-C. (Treatment: Prob > F = 0.2284) **f)** Heterogeneity of lattice spacing (σ_D_) of permeabilized psoas muscle at SL 2.4 and 3.0 before and after removal of fMyBP-C. (Treatment: Prob > F = 0.003) **g)** Intensity of the M3 reflection (I_M3_) of permeabilized psoas muscle at SL 2.4 and 3.0 before and after removal of fMyBP-C. (Treatment: Prob > F = 0.0001) **h)** T3 spacing (S_T3_) of permeabilized psoas muscle at pCa 6.0 at SL 2.4 and 3.0 before and after removal of fMyBP-C. (Treatment: Prob > F = 0.0377) **i)** Actin layer line 6 spacing (S_ALL6_) of permeabilized psoas muscle at SL 2.4 and 3.0 before and after removal of fMyBP-C. (Treatment: Prob > F = 0.0026) Statistics throughout are a repeated measure ANOVA design with SL, treatment, and their interaction as main effects, including a random effect on individual fiber. A Student’s t-test or Tukey’s HSD Post-Hoc test was performed on statistically significant differences and presented in figures as connecting letters. Dataset: (n=43 fibers, N=13 animals).

### fMyBP-C stabilizes lattice order during isometric contraction, reduces myosin head order, and shortens thin filament length during isometric contraction

Figure 6 shows a representative X-ray diffraction pattern collected from active SnoopC_2_ psoas muscle. We report on features of equatorial reflections, including the spacing of the 1,0 reflection (d10), the thick-filament only lattice plane spacing, and its width in equatorial space, which represents the degree of heterogeneity in the lattice and thick filament disorder (σ_D_)(40). Under activating conditions at pCa 6.0 (approximately the pCa_50_ for force generation), we found that lattice spacing decreased with increasing SL as expected. However, there were no significant differences in lattice spacing following removal of fMyBP-C (Fig. 6d). This result differs from the increase in lattice spacing following loss of fMyBP-C observed under relaxing conditions (23), but may be explained by the relatively greater influence of strongly bound cross-bridges to act on the thin filament and limit lattice expansion during Ca^2+^ activation. However, we found that the heterogeneity of lattice spacing (σ_D_) was significantly increased at both SL 2.4 and 3.0 following removal of fMyBP-C (Fig. 6f). The latter suggests a modest degradation in the lattice integrity during contraction when fMyBP-C is cleaved.

In order to further investigate the role of fMyBP-C in the orientation and order of myosin heads as well as the thin filament, we investigated the I_1,1_/I_1,0_ equatorial intensity ratio and M3, T3, M6 meridional reflections as well as the 6^th^ actin layer line (ALL6) in the X-ray diffraction patterns. The ratio of the 1,0 and 1,1 reflection intensities (I_1,1_/I_1,0_) reflects the distribution of mass between thick and thin filaments, where increases are typically associated with a shift of myosin heads to the ON state and/or binding of myosin heads to the thin filament. We found that a loss of fMyBP-C did not have a significant effect on the I_1,1_/I_1,0_ intensity ratio during contraction, suggesting that loss of fMyBP-C did not significantly affect the distribution of myosin heads away from the thick filament backbone or proportion of bound crossbridges under Ca^2+^ activated conditions. The I_1,1_/I_1,0_ intensity ratios at both short and long SL (SL 2.4 and 3.0) also remained unchanged during isometric contraction, as previously reported by others (41, 42). Next, the spacing of the M3 reflection (S_M3_), a measure of the axial periodicity of myosin heads, did not significantly change at either SL 2.4 or 3.0 following loss of fMyBP-C (Supplementary Figure x). Because S_M3_ is also taken as a measure of whether myosin heads are primarily in the OFF or ON state, results suggest that fMyBP-C does not contribute significantly to the OFF/ON distribution of myosin heads or numbers of bound myosin heads under active conditions, consistent with the I_1,1_/I_1,0_ intensity ratio. The intensity of the M3 reflection (I_M3_), however, decreased following removal of fMyBP-C (Figure 7g), consistent with an increase in the angular dispersion of myosin heads(43, 44), suggesting that loss of fMyBP-C promotes disorder of myosin heads. The spacings of other thick filament reflections, such as (S_M6_), indicative of thick filament length, were unchanged following removal of fMyBP-C (Supplementary Figure 4).

Finally, we sought to determine whether any thin filament structural changes occurred following fMyBP-C removal. We found that the spacing of the third-order troponin reflection (S_T3_) was significantly decreased following removal of fMyBP-C at both SL 2.4 and SL 3.0, suggesting a reordering of the troponin complex on the thin filament. Additionally, the spacing of actin layer line 6 (S_ALL6_), reflecting the pitch left-handed helix of actin, was significantly decreased following removal of fMyBP-C at both SL 2.4 and SL 3.0 µm (Fig. 6i).

## Discussion

### Myosin binding protein-C paralogs commonly influence Ca^2+^-sensitivity, cross-bridge cycling, and oscillatory contractions in striated muscles

In this study, we sought to identify key similarities and differences in the role of the three major MyBP-C paralogs in regulating muscle contractility, as well as identify key roles for fMyBP-C in the structural organization of isometrically contracting sarcomeres. Using three genetically modified mouse lines specific to slow (SpyC_1_), fast (SnoopC_2_), and cardiac-MyBP-C (SpyC_3_) that allow for selective removal of MyBP-C and replacement with recombinant MyBP-C in the respective muscle types, we were able to test mechanical properties of each muscle type in the presence and absence of MyBP-C. We found that selective cleavage of MyBP-C led to a reduction in Ca^2+^-sensitivity in both cardiac and fast twitch myocytes. Effects on Ca^2+^-sensitivity did not reach significance in soleus, a predominantly slow twitch muscle, likely due to significant co-expression of fast and slow MyBP-C (since fMyBP-C would not be cleaved in skeletal muscles from SpyC_1_ mice). However, recombinant slow MyBP-C was able to restore Ca^2+^-sensitivity when added to SpyC_3_ cardiac myocytes (Figure 4f), so all 3 paralogs apparently share the common ability to shift the tension-pCa relationship to the left. Similarly, cleavage of each MyBP-C paralog increased cross-bridge cycling rates at submaximal Ca^2+^ activation as measured by *k_tr_*. This result is in good agreement with previously reported changes in both cardiac and fast skeletal MyBP-C KO mice as well as previous work with the SnoopC_2_ and SpyC_3_ mouse lines(22, 45, 46). Moreover, loss of each MyBP-C paralog led to the appearance of spontaneous oscillatory contractions at sub-maximal Ca^2+^ activations near the pCa_50_ for force generation. While the precise mechanisms by which oscillatory contractions are induced and propagated are still unclear, the phenomenon is an intrinsic, emergent property of sarcomeres that has been likened to a third state of muscle activity between activation and relaxation(29, 47). Notably, because force oscillations occur at Ca^2+^ concentrations expected to be encountered in living muscle and are influenced by metabolites such as ADP/P_i_(48), tension instabilities are likely to have physiological relevance. Consistent with this idea, mutations in slow MyBP-C lead to force oscillations that are associated with skeletal muscle tremor in humans(17, 49, 50). Results presented here therefore suggest that a novel functional role of all 3 MyBP-C paralogs is to limit mechanical force instabilities and promote contractile synchrony. Conversely, loss of MyBP-C or mutations in MyBP-C underlie contractile dysfunction in different muscle types (51).

### Myosin Binding Protein C paralogs differentially contribute to tension transients in striated muscle

While we identified several common functional effects of the different MyBP-C paralogs as discussed above, a notable departure from this trend was evident when comparing the differential responses to a rapid 2% stretch in each muscle type which we found were attributable to expression of the different MyBP-C paralogs. For instance, under control conditions all 3 muscle types (cardiac, fast and slow skeletal) had unique responses to rapid stretch that were mostly stereotypical for that muscle type as previously described by others(19, 33, 34, 52, 53). This was most evident when comparing the responses of cardiac and psoas (fast skeletal) muscles, where cardiac muscle exhibited a canonical 3 phase response that included a final phase such that after reaching a nadir (P_2_), steady state force rises above the pre-stretch force, P_3_. (SA, discussed in more detail below). By contrast, P_2_ was highly elevated in psoas muscle and the phase 3 rise in force was typically minimal and/or short-lived. Notably, classical SA behavior is generally considered minimal or absent in fast twitch muscle, although enhancement of SA in skeletal muscles may be important under conditions of fatigue (e.g., high inorganic P_i_) (34, 52, 54). Strikingly, as shown in Figure 3, we found that loss of fMyBP-C after TEVp cleavage in psoas muscle completely altered the stereotypical response to stretch and completely converted the response into one resembling that of cardiac muscle, including a significant decrease in P_2_ and the appearance of a prominent phase 3 SA (delayed force increase) response. Conversely, when fMyBP-C was added to cardiac myocytes (Fig. 5), the tension transient converted into a response more closely resembling one typically found in fast skeletal muscle where P_2_ was elevated and phase 3 was eliminated. Taken together, our results provide compelling evidence that the different MyBP-C paralogs contribute significantly to the stereotypical tension transient responses in their respective muscle types. Furthermore, to our knowledge, this is the first report showing that loss of fMyBP-C confers classic SA-like properties to fast twitch skeletal muscle fibers.

Tension transient responses to stretch, including SA responses, are thought to be mediated largely by changes in cross-bridge cycling kinetics (30). Accordingly, numerous factors that are known to impact cross-bridge dynamics, including calcium, ADP, and P_i_ concentration, the extent of stretch, temperature, and the contribution of other sarcomere proteins, such as thin filament proteins as well as myosin heavy chain isoforms, have all have been shown to significantly impact the shape and rate of tension transient responses to stretch(34, 54–56). In this respect, it may not be surprising that MyBP-C, known to affect cross-bridge cycling kinetics, also influences tension transients. Indeed, consistent with results reported here, Stelzer et al (2006) reported increased K*_rel_*, reduced P_2_, and increased *K*_df_ rates in cMyBP-C KO mice, effects that were largely replicated by PKA phosphorylation of WT cMyBP-C (33, 39). PKA phosphorylation of sMyBP-C in slow twitch fibers also increased force development rates (57). Our results thus confirm and extend previous findings by showing that all 3 MyBP-C paralogs limit cross-bridge cycling kinetics in their respective muscle types. However, to further explore the extent to which each paralog by itself was sufficient to determine the overall shape of tension transients when expressed in a similar background, we also utilized the SpyC_3_ cardiac mouse line that primarily expresses a single myosin isoform (ɑ-myosin) in cardiac myocytes to systematically replace endogenous cMyBP-C with recombinant cardiac, slow, and fast MyBP-C. Again, we found that each MyBP-C paralog conferred a characteristic tension transient in cardiac muscle, thus allowing us to attribute observed changes in the tension transients primarily to the different paralogs of MyBP-C rather than to different myosin isoforms or other confounding variables.

### MyBP-C paralogs determine the extent of strain-induced cross-bridge detachment, P_2_

A key difference that emerged when comparing the tension transients in the different muscle types was the extent of the P_2_ nadir, which presumably reflects the magnitude of cross-bridge detachment in response to strain in each muscle type. Notably, P_2_ was reduced following loss of MyBP-C in all 3 muscle types, suggesting that a normal function of MyBP-C is to limit strain-induced cross-bridge detachment. In cardiac muscle, strain-dependent cross-bridge detachment is emerging as a key regulator of the mechanical control of relaxation where stretch enhances LV relaxation during the transition from late systole to early diastole(55, 58, 59). In fact, Stehle et al. concluded that load-dependent cross-bridge detachment is the only mechanism that explains how cardiac muscle can relax as quickly as it contracts(60). Consistent with this, small stretches during the slow phase of myofibril relaxation were shown to accelerate the onset of sarcomere “give” which initiates the fast exponential phase of relaxation(61). We also recently showed that loss of cMyBP-C significantly accelerates the fast phase of relaxation suggesting that cMyBP-C provides an internal load (potentially by cross-linking thick and thin filaments) or otherwise stabilizes sarcomere dynamics to slow relaxation(24).

In fast skeletal muscle acceleration of strain-dependent cross-bridge detachment after loss of MyBP-C may contribute to the appearance of SA-like properties. For instance, Wakefield et al. modeled cardiac tension transients in terms of a model where the shape of the transient was the sum of 3 main factors: a passive Power-law relaxation term, a Reversal term that is responsible for the transient drop in force (P_2_ nadir), and a Drag term that represents the initial fast rise in tension and relaxation immediately after the stretch(59). If the Reversal term in skeletal muscle is small as suggested by Davis and Epstein(32), and further limited by MyBP-C, then the shape of the tension transient in skeletal muscle would be expected to be dominated by a combination of the Drag and Power-law terms. However, if loss of fMyBP-C increases strain-dependent cross-bridge detachment (*g*_1_ in their model), then the Reversal term would become greater and deepen the amplitude of the P_2_ nadir(59). At the same time, loss of MyBP-C could further reduce the Drag term as well as leading to an overall increase in the R/D ratio. Together, the two effects would lead to the appearance of an SA response as observed following the loss of fMyBP-C.

Through which interactions are the various MyBP-C isoforms able to affect the Drag and Reversal terms? If MyBP-C links the thick and thin filaments and is able to slow filament sliding(45, 57), then this interaction would be expected to increase the Drag term. MyBP-C’s presence may also limit the Reversal term, for instance, by preventing P_i_-dependent reversal of the powerstroke. Consistent with this cMyBP-C slows phosphate (P_i_)-dependent myosin cross-bridge detachment and prolongs cross-bridge lifetimes in cardiac muscle (62, 63). Tanner et al. further concluded that cMyBP-C inhibits P_i_-dependent reversal of the powerstroke or otherwise stabilizes cross-bridge attachment in a way that increases the probability of myosin completing its powerstroke(64). Notably, to account for differences in relative rates of P_2_ (P_2-slow_) in different fiber types, Davis and Epstein argued that P_i_-dependent reversal of the powerstroke is largely absent in fast fibers, whereas it is prominent in slow fibers(32). Straight et al. also concluded that a reverse stroke of slow skeletal myosin accounts for augmentation of SA in slow skeletal muscle, but not in fast skeletal muscle(34), although the same group later found that increases in [P_i_] concentration, such as during muscle fatigue, augments SA even in fast twitch fibers(54). Using optical tweezers and synthetic cardiac myofilaments, Hwang et al. demonstrated that a reversal stroke is present in cardiac myosin, but absent in fast skeletal myosin, where it is a key determinant of cardiac contraction and relaxation and is further necessary to produce stable force even at low levels of Ca^2+^ activation(65). While the reversal stroke (or absence of) is therefore likely intrinsic to the myosin isoform, our results suggest that MyBP-C paralogs are key regulators of cross-bridge detachment rates, potentially via modulation of P_i_-dependent reversal of the powerstroke(66).

SA is a phenomenon best described in some insect flight muscles that operate at high frequencies (100-1000 Hz) where muscle is turned on and off in response to mechanical signals rather than being driven by slower and more energetically costly chemical and electrical signals(35, 67, 68). In these so-called asynchronous muscles, a single action potential elicits a sustained rise in [Ca^2+^] that is permissive for multiple cycles of muscle activation and relaxation driven by muscle stretch. The initiating mechanical event, muscle stretch, itself is the result of contraction of an antagonistic muscle such that asynchronous muscles act in pairs. By accelerating strain-induced cross-bridge detachment, it is plausible that stretch synchronizes relaxation of a population of cross-bridges that contract together resulting in SA. In this respect it is notable that MyBP-C, which appears to limit strain-induced cross-bridge detachment as we show here, is absent in insect flight muscles, although other proteins such as projectin and flightin may perform analogous roles(69–71). While mammalian cardiac and skeletal muscles utilize a synchronous contractile mechanism (where a single action potential leads to a single twitch), SA occurs to some extent in both muscle types and may play important physiological roles. Examples include coordinating proper ejection timing of the ventricle and mechanical control of relaxation in the heart (55, 58, 72), or during exercise where contraction of one muscle may lead to stretch in its antagonistic partner, priming it for contraction moments later(52, 54). Our results suggest that by modulating strain-dependent cross-bridge detachment MyBP-C fine-tunes the contribution of SA in different muscle types and that regulation of MyBP-C, for instance by phosphorylation within a given muscle type, may thus contribute to a greater or lesser dependence on mechanical signals for activation and relaxation(33, 57).

### Potential mechanisms for MyBP-C regulation of the tension transient response

To gain additional insights into mechanisms by which loss of MyBP-C affects cross-bridge kinetics, we investigated structural effects of loss of MyBP-C using X-ray diffraction analyses of Ca^2+^-activated psoas muscle before and after cleavage of MyBP-C with TEVp. We chose psoas muscle for X-ray diffraction studies because the effects of loss of MyBP-C were greatest in psoas muscle (e.g., the reduction in P_2_ was greatest) and because we had previously characterized loss of MyBP-C in psoas muscle under relaxed conditions(23). In that study we found that loss of MyBP-C expanded the sarcomere lattice, reduced the length of the thin (actin) filament, and repositioned myosin heads from their OFF to ON positions on the thick filament. These results were overall consistent with effects of MyBP-C to stabilize the inhibited state of myosin and limit lattice expansion potentially through interactions of MyBP-C with actin. By contrast, here we found under Ca^2+^-activated conditions that loss of MyBP-C did not significantly affect lattice spacing possibly because strongly bound cross-bridges effectively limit the separation of the thick and thin filaments. However, an indicator of lattice disorder or heterogeneity, σ_D_, was significantly increased at both SL 2.4 and SL 3.0 µm following loss of MyBP-C. Palmer et al., also reported greater myofilament disorder in a mouse model of cMyBP-C knockdown(63). σ_D_ is increased during active conditions compared to passive conditions, likely due to contributions of cross-bridge cycling that tend to move the positions of both thick and thin filament relative to their equilibrium position in the lattice(73). Therefore, results suggest that MyBP-C appears to plays a role in maintaining thick and thin filaments near their equilibrium positions during contraction, possibly by linking thick and thin filaments together. If so, then MyBP-C may promote efficient/strong cross-bridge binding by maintaining alignment of binding sites relative to one another on thick and thin filaments. Conversely, an increase in lattice disorder with loss of MyBP-C may lead to less efficient binding of myosin cross-bridges and/or promote cross-bridge detachment, thereby decreasing myosin duty ratio and increasing cycling rates(74, 75). The idea that MyBP-C links thick and thin filaments together is also consistent with MyBP-C acting as a drag force or internal load that slows cross-bridge cycling kinetics and relaxation(24, 45, 57).

MyBP-C also had significant effects on both the myosin head position and the thin filament. The proportion of myosin heads in the ON state appeared unchanged following loss of MyBP-C as indicated by the spacing of the M3 reflection (S_M3_)(Supplementary Fig. X3), but the ordering of the myosin heads relative to the thick filament backbone was significantly decreased at both SLs as indicated by a reduction in the M3 intensity (I_M3_) (Fig. 6). Increased disorder of the myosin heads captured in the M3 intensity may be indicative of less efficient cross-bridge binding to the thin filament and may explain the faster cycling rates identified in mechanics experiments. Finally, loss of MyBP-C decreased both the spacing of the third order troponin reflection (S_T3_) and the spacing of actin layer line 6 reflection (S_ALL6_). The reduced spacing of these two actin-based reflections indicate an overall reduction of thin filament length, consistent with the ability of MyBP-C to bind to and exert a strain on the thin filament. In line with this, MyBP-C is essential for thick filament stability and lattice rigidity(62, 63, 76). Notably, filament compliance can affect stability of strongly bound cross-bridges and relaxation rates(77–79). Taken together, the changes in X-ray diffraction reflections are consistent with a role for MyBP-C in minimizing overall disorder of the sarcomere lattice during contraction and maintaining myosin heads in ordered positions. Whereas proper alignment of thick and thin filaments relative to one another may promote strong cross-bridge binding and cross-bridge stability, loss of thick and thin filament alignment may promote cross-bridge detachment, especially under strain.

## Limitations

A limitation of our X-ray diffraction analysis was that structural changes were investigated during isometric contraction, making it difficult to draw conclusions about whether dynamic changes occur during rapid stretch that are not captured during steady state Ca^2+^ activation. Future studies will include time-resolved X-ray diffraction measurements taken as the muscle is being stretched in order to obtain structural information during the various phases of the tension transient. Another limitation was that the soleus muscle contains a mixed population of fast type and slow type fibers and we did not attempt to differentiate their different contributions in mechanical force measurements. As a result, the mechanical contribution of slow MyBP-C was likely underestimated due to the significant presence of fast MyBP-C which was not subject to cleavage in SpyC_1_ mice. In addition, there are at least two splice variants of slow skeletal MyBP-C expressed in mouse and we did not attempt to determine their relative expression(21).

## Conclusions

Major conclusions from this study are that all 3 paralogs of MyBP-C (cardiac, fast skeletal and slow skeletal) exert robust functional effects on force generation in their respective muscle types. Using selective cleavage of each paralog we showed that all 3 had common functional effects to increase Ca^2+^ sensitivity of tension and limit cross-bridge cycling rates at submaximal levels of activation. In addition, all 3 were effective at damping force oscillations that appeared following loss of MyBP-C when MyBP-C was cleaved with TEVp. Finally, all 3 paralogs slowed strain-dependent cross-bridge detachment rates which was evident as accelerated detachment rates (*K*_rel_) and a lower force nadir (P_2_) following loss of MyBP-C when muscles were subjected to a rapid stretch. However, the overall transient force response to a rapid stretch was unique in each muscle type and dependent on the MyBP-C paralog expressed. In particular, MyBP-C paralogs uniquely modulated strain dependence of cross-bridge detachment and thereby influenced SA properties in each muscle type. This regulation of cross-bridge relaxation may have significant physiological importance, as strain-induced cross-bridge shortening is key to rapid relaxation at onset of diastole to allow for proper ventricular filling, and may also enable rapid bursts in contraction in skeletal muscle.

## Materials and Methods

### Animal procedures

All animal procedures were performed under protocols approved by the Institutional Animal Care and Use Committee (IACUC) at the University of Arizona. SpyC_1_, SnoopC_2_, and SpyC_3_ (C57BL6/NJ) mice were created by the Genetically Engineered Mouse Models (GEMM) core at the University of Arizona. All mice were bred and housed in a pathogen-free barrier animal facility at the University of Arizona. Animal genotypes were confirmed using PCR of DNA extracted from toe-clip samples and amplicons visualized on 1% agarose gels.

Mice used in this study were euthanized via cervical dislocation following anesthesia with isoflurane. Left ventricles, psoas, and soleus muscles were immediately dissected and flash frozen in liquid nitrogen for long-term storage at −80°C for force measurements. For X-ray diffraction measurements, muscles were permeabilized using glyercol (1:1 rigor:glycerol where Rigor solution contained (in mM): KCl (100), MgCl_2_ (2), ethyleneglycol-bis(β-aminoethyl)-N,N,N’,N’-tetraacetic acid (EGTA, 5), Tris (10), dithiothreitol (DTT, 1), 1:100 HALT™ Protease Inhibitor Cocktail (ThermoFisher Scientific, Waltham, MA, USA)) at 4°C for 48 hours. After 48 hours the solution was refreshed and then muscles were stored up to 60 days at −20°C until use.

### Recombinant protein expression and purification

Protein plasmids with desired DNA sequences were purchased from Genscript (Piscataway, NJ, USA). BL21 and DE3 competent cells (C2527, New England Biolabs, Ipswich, MA, USA) were transfected with plasmid DNA and protein purification was performed as described previously.(13, 22)

### Muscle force measurements

Tissue was thawed in a skinning solution consisting of a Relax Buffer containing 1% Triton X-100, 0.1% Saponin, and 1:100 HALT™ Protease Inhibitor Cocktail (ThermoFisher Scientific, Waltham, MA, USA). Relax buffer contained (in mM): 100 KCl, 10 imidazole, 2 EGTA, 5 MgCl_2_, 4 ATP (A2383, Sigma, St. Louis, MO, USA), and 1:100 HALT™ Protease Inhibitor Cocktail at pH of 7.0. Multicellular preparations of cardiac cells and skeletal muscle fibers were then isolated by mechanical disruption using a homogenizer (Omni International 60CE54. Model: TH115). After homogenization, cardiac and skeletal muscle cells were incubated with rotation in a skinning solution at 4°C for 15 and 30 min, respectively. Detergent-permeabilized cardiac and skeletal muscle cells were then washed thoroughly with fresh Relax buffer to remove residual detergents. Next, cardiac cells and skeletal muscle fibers (∼80-200 mm in length, 50-100 mm in width) were selected and attached to a high-speed positional motor (Model: 315C-I, Aurora Scientific Inc., Aurora, Ontario, Canada) and a force transducer (Model 403 A series, Aurora Scientific Inc., Aurora, Ontario, Canada) using aquarium sealant (Marineland, 100% clear silicone rubber or Loctite, Clear Silicone Waterproof Multipurpose Adhesive Sealant). The positional motor and force transducer were attached over a temperature-adjustable 8-well platform (Model 803B, Aurora Scientific Inc., Aurora, Ontario, Canada) regulated by a thermocouple (825A, Aurora Scientific Inc., Aurora, ON, Canada). The 8-well platform was then mounted on the stage of an inverted light microscope (Model IX-53, Olympus Instrument Co., Japan). The aquarium sealant was allowed to cure for 30 min before the experiment was started. SL was then measured using video imaging in passive conditions (at pCa 9.0) and SL was set to a final length of 2.3 μm and 2.4 μm for cardiac and skeletal muscle, respectively. All muscle force measurements were obtained at 15°C using a range of pCa solutions with free Ca^2+^ pCa 9.0 to 4.5, where pCa = -log[Ca^2+^]. Force measurements (F) recorded at each submaximal pCa solution were normalized to the maximal force (F_0_) at pCa 4.5 and to cross-sectional area of the muscle fiber assuming circular dimensions. Data was fitted using a sigmoidal four-parameter logistic curve within Graphpad Prism. *k_tr_* was assessed by performing a slack-restretch test, where the muscle was shortened to 80% of its reference length for 20 ms, stretched to 105% reference length for 5 ms and returned to reference length. Tension transient parameters were assessed by performing a rapid 2% stretch over 5 ms and held for 3 s. Cross-bridge cycling rates (*k_tr_* and shortening induced *k_df_*) and cross-bridge relaxation (*k*_rel_), were fit using a single exponential curve and normalized to maximal rate (*k_0_*) at pCa 4.5 and plotted against isometric force.

### N-terminal MyBP-C cleavage and *in situ* paste of recombinant proteins

Mechanics experiments typically consisted of 3 consecutive sets of measurements: First, before N-terminal cleavage of MyBP-C (control), second, after N-terminal cleavage of MyBP-C with TEVp, and third in a subset of experiments after *in situ* paste of recombinant MyBP-C. Proteolytic cleavage of domains C0-C7 in cardiac cells and C1-C7 in skeletal muscle cells was performed by incubating myocytes in 20 μM AcTEV™ Protease (ThermoFisher, 12575015) at 25°C for 30 min while the permeabilized muscle preparation was attached to the high-speed positional motor and force transducer. After TEVp incubation, cells were rigorously washed in fresh relax buffer (3 x for 3 min) to remove TEV protease as well as cleaved endogenous MyBP-C N-terminal domains. Paste of recombinant protein was done by incubating permeabilized cell preparations for 15 min at 15°C with 20 µM recombinant protein of choice containing a SpyCatcher sequence. Myocytes were then washed (3 x for 3 min) using fresh relax buffer to remove excess (non-ligated) recombinant protein. N-terminal MyBP-C cleavage and protein ligation was assessed using SDS-PAGE gel electrophoresis and western blot as described previously(23).

### Small angle X-ray diffraction

All X-ray diffraction experiments were performed with the small-angle instrument at the BioCAT beamline 18ID at the Advanced Photon Source, Argonne National Laboratory. Muscle experiments were performed both before and after the facility wide upgrade of the Advanced Photon Source with changes to beam flux (Pre: ∼3×10^12^, Post: ∼5×10^13^ photons per second). The X-ray diffraction beam (0.103 nm wavelength), focusing at the detector plane (Pre: ∼0.06 x 0.15 mm. Post: 0.02 x 0.02 mm), sample to detector distance (∼2m), and detector (Mar 165, Rayonix Inc., USA) are identical to our previous study in relaxed muscle fibers(23). An in-line camera was present to properly target beam onto muscle fibers and assess for any irregularities in attachment of the muscle. Muscle fibers were attached to custom fabricated clamps and connected to a high-speed length controller (322C, Aurora Scientific, Canada) and a force transducer (402A, Aurora Scientific, Canada) with a sampling rate of 1,000 Hz. Measurements from the force transducer to identify activation levels of the fibers were visualized using Real Time Muscle Data Acquisition and Analysis system (600A, Aurora Scientific, Canada). Experiments were performed at ∼25°C. SL was set at SL 2.4 µm or SL 3.0 µm using a 4-mW Helium-Neon laser based on width of the first-order diffraction. Muscles were activated by removal of relaxing solution (in mM: potassium propionate (45.3), N,N-Bis(2-hydroxyethyl)-2-aminoethanesulfonic acid BES (40); EGTA (10), MgCl_2_ (6.3), Na-ATP (6.1), DTT (10), protease inhibitors [Complete], pH 7.0), three 30 second incubations with pre-activating solution (in mM: potassium propionate (45.3), N,N-Bis(2-hydroxyethyl)-2-aminoethanesulfonic acid BES (40); MgCl_2_ (6.3), Na-ATP (6.1), DTT (10), protease inhibitors [Complete], pH 7.0), and finally by incubation with pCa 6.0 solution (in mM: potassium propionate (170), magnesium acetate (2.5), MOPS (10) and ATP (2.5), and CaEGTA and K_2_EGTA were mixed at different proportions to obtain different pCa (−log[Ca^2+^]), pH 7.0) values. Following incubation with pCa 6.0, tension was monitored until the muscle reached a steady-state level of force, upon which the bath was lowered with a servo-motor and X-ray images were taken in open air. No part of the sample was exposed to the X-ray beam more than once. Samples then underwent TEVp protease incubation as described above, and the process was repeated.

### X-ray Diffraction Analysis

Analysis of x-ray diffraction patterns was performed using MuscleX (version 1.22.0) open-source data reduction and analysis package (BioCAT, Argonne National Laboratories) as previously described(23, 80).

### Statistics

Statistical analyses were performed either in Graphpad Prism or JMP Pro (V16, SAS Institute Inc., Cary, NC, USA). All measurements reported as the mean ± SEM. Repeated measures one-way ANOVA tests with main effects pCa, TEVp treatment, and their interaction, as well as an individual fiber random effect were performed on measurements before TEV protease cleavage, after TEV protease cleavage, and when present, after addition of recombinant protein. A mixed effects test was used for any missing values. A paired t-test was used specifically to compare pCa50 values. The significance level was p<0.05. For additional statistical details see tables contained in Supplemental Dataset 1.

## Supporting information

Supplemental Video 2 - Soleus SPOC Video

Supplemental Dataset 1 - Statistical Tables

Supplemental Video 1 - Psoas SPOC Video

## Acknowledgments

This research was performed on APS beam time award(s) from the Advanced Photon Source, a U.S. Department of Energy (DOE) Office of Science user facility operated for the DOE Office of Science by Argonne National Laboratory under Contract No. DE-AC02-06CH11357. BioCAT is supported by grant P30 GM138395 from the National Institute of General Medical Sciences of the National Institutes of Health. The content is solely the responsibility of the authors and does not necessarily reflect the official views of the National Institute of General Medical Sciences or the National Institutes of Health. We would like to thank Dr. Thomas C Irving for expertise and help with editing the manuscript and John Sullivan for generating a video of SPOC in soleus muscle. Funding for this study was provided by the National Institute of Health (HL080367 [SPH]), HL140925 [SPH], AR081935 [SPH], T32 HL007249 [NME]), the American Heart Association (827628 [NME]), and German Research Foundation (DFG) 454867250 to ALH.

## Author Contributions

N.M.E, A.L.H, and S.P.H conceptualized the project, , N.M.E conducted the investigation, N.M.E visualized the datasets, S.P.H was the project administrator, A.L.H, and S.P.H supervised the study, N.M.E wrote the original draft, all authors reviewed, edited, and agreed to the final draft.

