## Supplemental Appendix for "Myosin binding protein-C limits strain induced cross-bridge detachment in response to rapid stretch in cardiac and skeletal muscle"

* Samantha P. Harris

Samantha P Harris*


**This PDF file includes:**

Figures S1 to S4
Legends for Movies S1 to S2

Dataset S1

**Other supporting materials for this manuscript include the following:**

Movies S1 to S2

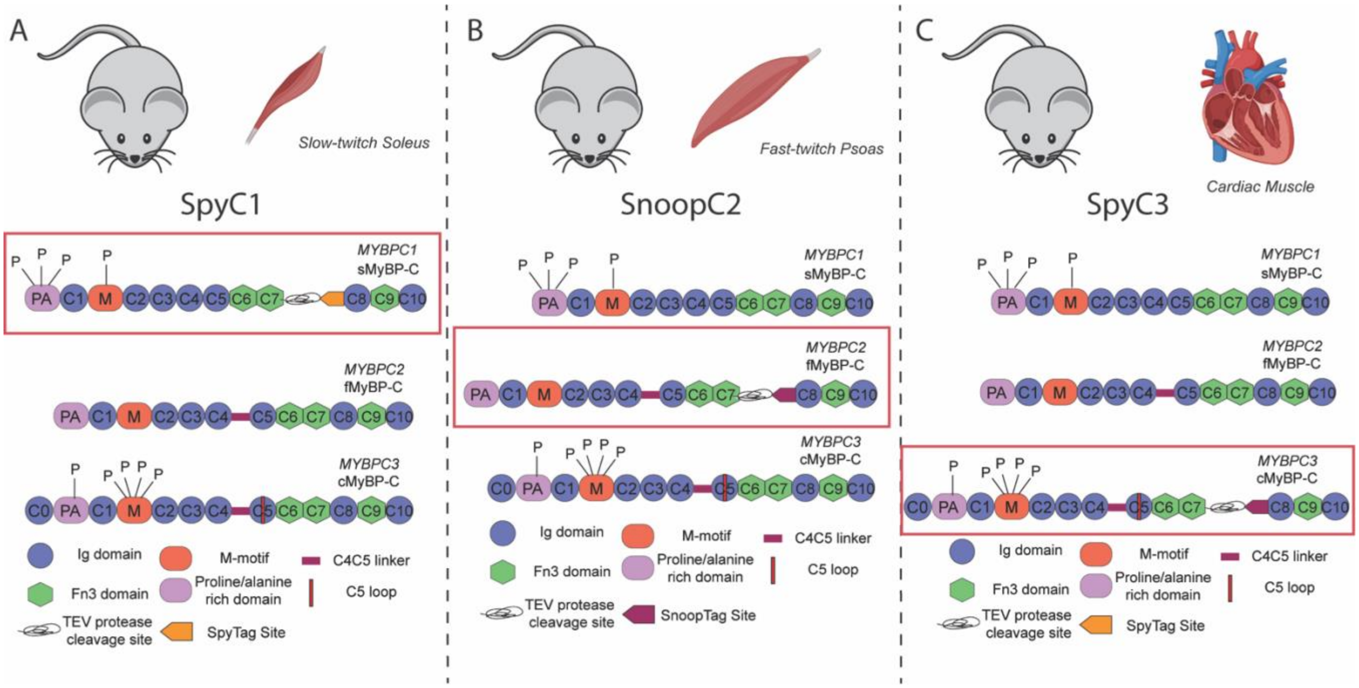


**Figure S1: Detailed description of SpyC1, SnoopC2, and SpyC3 mice. A**: A graphical description of the SpyC1 mouse, where the *Mybpc1* encoded slow-twitch myosin binding protein-C (sMyBP-C) contains a TEV protease recognition site and a SpyTag site between domains C7 and C8, while *Mybpc2* encoded fast-twitch MyBP-C (fMyBP-C) and *Mybpc3* encoded cardiac MyBP-C (cMyBP-C) contain no modifications. **B:** A graphical description of the SnoopC2 mouse, where the *Mybpc2* encoded fMyBP-C contains a TEV protease recognition site and a SnoopTag site between domains C7 and C8, while *Mybpc1* encoded sMyBP-C and *Mybpc3* encoded cMyBP-C contain no modifications. Specific domain names and paralog unique features are outlined below the three protein illustrations. **C:** A graphical description of the SpyC3 mouse, where the *Mybpc3* encoded cMyBP-C contains a TEV protease recognition site and a SpyTag site between domains C7 and C8, while *Mybpc1* encoded sMyBP-C and *Mybpc2* encoded fMyBP-C contain no modifications. Specific domain names and paralog unique features are outlined below the three protein illustrations. Protein Kinase A (PKA) mediated phosphorylation sites are indicated by “P”.

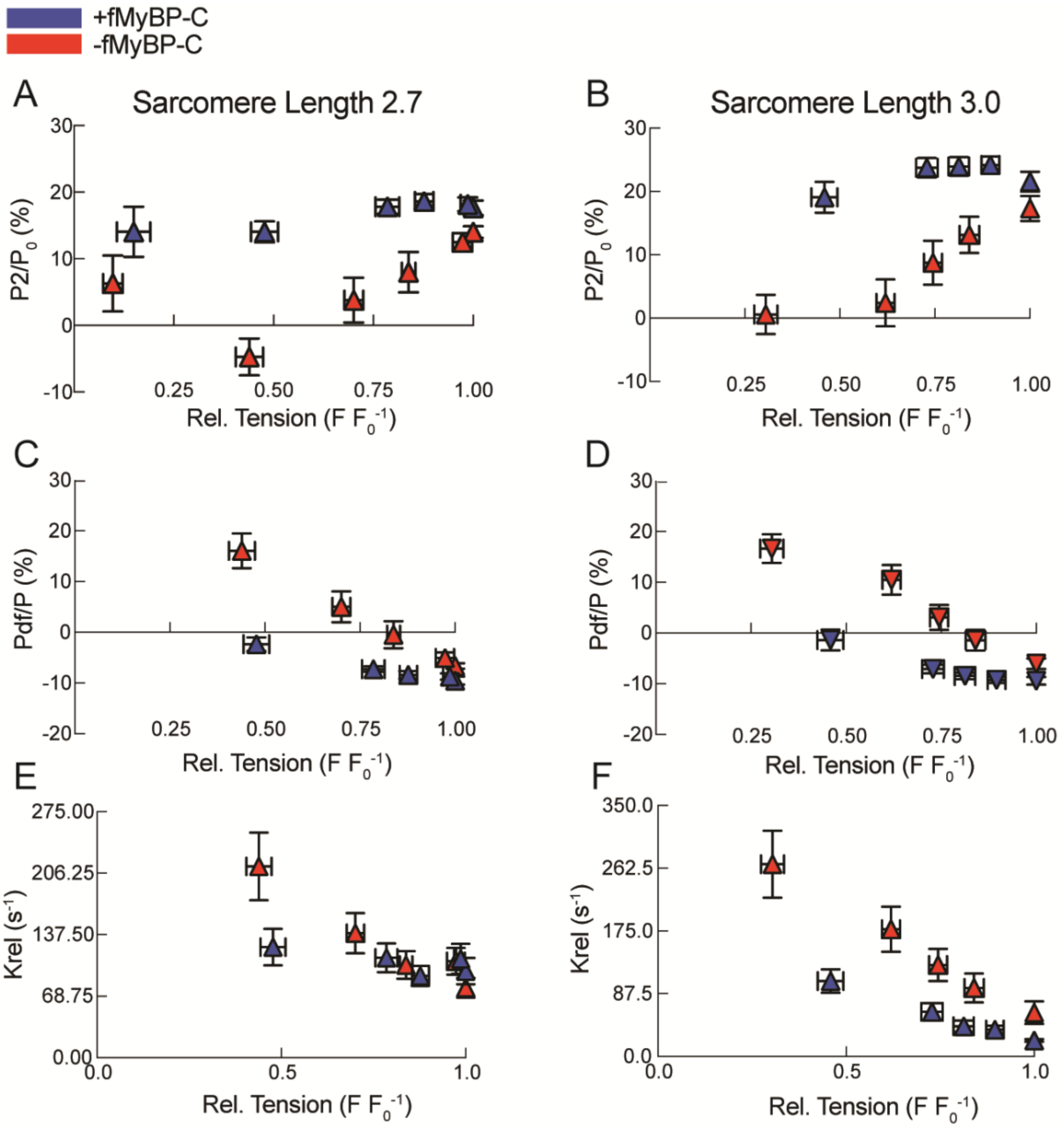


**Figure S2: Removal of fMyBP-C results in decreased P2 and increased Pdf at sarcomere lengths 2.7 and 3.0. A-B:** P2 as a percentage pre-stretch steady state force ($P_0$) before (blue) and after (red) removal of fMyBP-C in fast-twitch psoas muscle at SL 2.7 (A) and SL 3.0 (B). In both cases, P2 drops to or below $P_0$ at $pCa_{50}$ values following removal of fMyBP-C. (SL 2.7: Treatment*pCa: Prob > F = 0.0366. SL 3.0: Treatment*pCa: Prob > F = 0.0008) Connecting letters report shows significant differences **C-D:** Pdf as a percentage of $P_0$ before (blue) and after (red) removal of fMyBP-C in fast-twitch psoas muscle at SL 2.7 (C) and SL 3.0 (D). Pdf shifts into positive values between 50-75% of maximal activation at both SL 2.7 and SL 3.0 (SL 2.7: Treatment*pCa: Prob > F = 0.0008. SL 3.0: Treatment*pCa: Prob > F = 0.0002). **E-F:** Crossbridge relaxation rate following rapid stretch ($K_{rel}$) before (blue) and after (red) removal of fMyBP-C in fast-twitch psoas muscle at SL 2.7 (E) and SL 3.0 (F). Krel increased at half-maximal values in both SL 2.7 and SL 3.0 (SL 2.7: Treatment*pCa: Prob > F = 0.0054. SL 3.0: Treatment*pCa: Prob > F = 0.0073). Statistics throughout are repeated measures ANOVA design with pCa, treatment, and their interaction as main effects. A random effect was assigned to individual fibers. A paired t-test or Tukey's Honestly Significant Difference post-hoc test was performed on significant main effects.

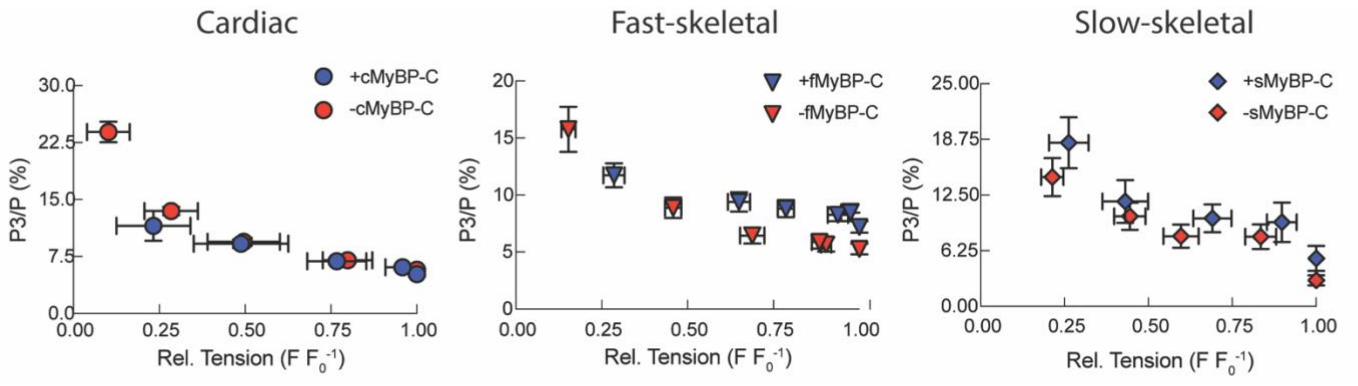


**Figure S3: Percent increase of the post-stretch steady-state in cardiac, fast-twitch psoas, and slow-twitch soleus muscles. A:** Quantification of the post-stretch steady-state force (P3) as a percentage of pre-stretch steady-state force in the presence (blue circle) and absence (red circle) of cardiac myosin binding protein-C (MyBP-C). While a significant interaction effect was found, P3 at values of similar levels of activation were not different from one another. (Treatment*pCa: Prob > F = 0.0001) **B:** Quantification of P3 as a percentage of pre-stretch steady-state force in the presence (blue triangle) and absence (red triangle) of fast-skeletal MyBP-C (Treatment*pCa: Prob > F = 0.0027) **C:** Quantification of P3 as a percentage of pre-stretch steady-state force in the presence (blue diamond) and absence (red diamond) of slow-skeletal MyBP-C (Treatment: Prob > F = 0.1492). Statistics throughout are a repeated measures ANOVA design with pCa, treatment, and their interaction as main effects. An individual fiber random effect was also included. Further statistical information can be found in Table/Dataset XXX. Dataset: (Cardiac: n=13, N=4. Psoas: n=10, N=3. Soleus: n=8, N=3 +sMyBP-C, N=4 -sMyBP-C).

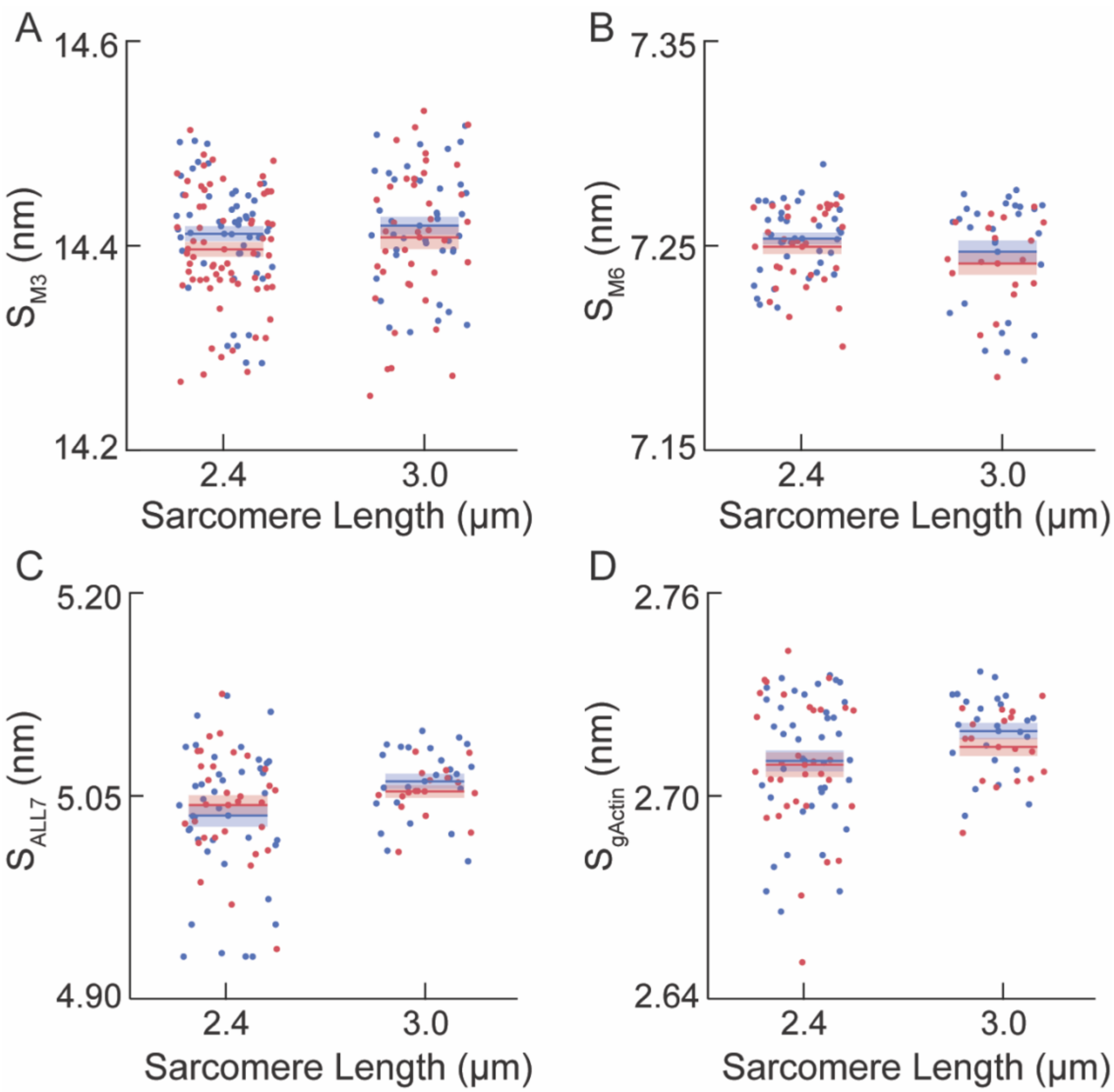


**Figure S4. M3, M6, ALL7, and gActin spacing remain unchanged following removal of fMyBP-C in half-maximally activated fast-twitch psoas muscle. A:** Spacing of the M3 meridional reflection at SL 2.4 and SL 3.0 in fast-twitch psoas muscle before (blue) and after (red) removal of fMyBP-C. (Treatment Prob > F = 0.1249) **B:** Spacing of the M6 meridional reflection at SL 2.4 and SL 3.0 in fast-twitch psoas muscle before (blue) and after (red) removal of fMyBP-C. (Treatment: Prob > F = 0.2574) **C:** Spacing of the 7$^{th}$ actin layer line (ALL7) meridional reflection at SL 2.4 and SL 3.0 in fast-twitch psoas muscle before (blue) and after (red) removal of fMyBP-C. (SL: Prob > F = 0.0308. Treatment: Prob > F = 0.9459) **D:** Spacing of gActin monomers at SL 2.4 and SL 3.0 in fast-twitch psoas muscle before (blue) and after (red) removal of fMyBP-C. (SL: Prob > F = 0.0439. Treatment: Prob > F = 03951). Data are presented as mean + s.e.m. Statistics throughout are a repeated measure ANOVA design with SL, Treatment, and the interaction as main effects, including a random effect on individual fibers. Dataset: (n=43 fibers, N=13 animals).

**Dataset S1 (separate file):** An excel spreadsheet containing tables indicating the specific statistical tests, means and standard errors, and statistical significances of each graph in the main text.

**Movie S1 (separate file):** Representative video showing the spontaneous oscillatory contractions (SPOC) that occur following removal of fMyBP-C in fast-twitch psoas muscle SPOC occurs during activation and persist once the muscle has reached a steady-state force. SPOC consistently occurs in a range of activation between 35-70% of maximal activation.

**Movie S2 (separate file).** Representative video showing the spontaneous oscillatory contractions that occur following removal of sMyBP-C in slow-twitch soleus muscle. SPOC begins during activation and persists once the muscle has reached a steady-state force. SPOC consistently occurs in a range of activation between 35-70% of maximal activation.
